## Supplementary Figures for "Suppression without inhibition: How retinal computation contributes to saccadic suppression"

#### Figure S1

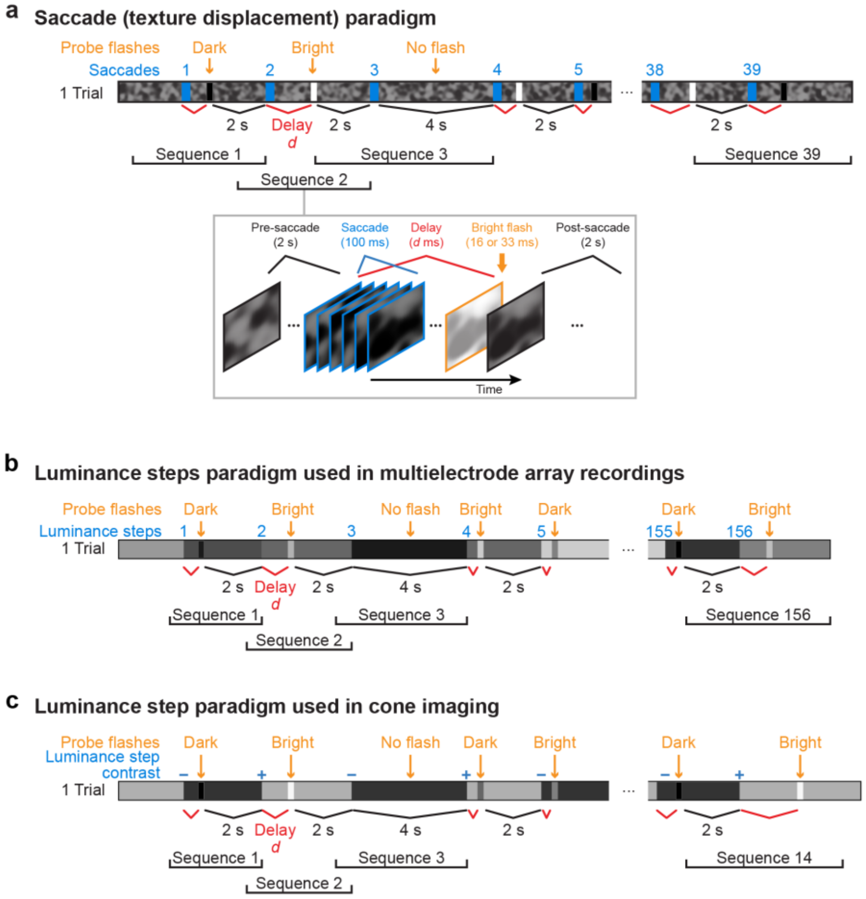


**Figure S1 Stimulation paradigms.**

**a.** Example trial of the temporal sequence of saccades and probe flashes. A trial started with turning on the texture over the retina. To mimic saccades (blue windows), the texture was displaced for 6 consecutive frames (100 ms) to a new position relative to the retina. After a certain delay *d*, a dark or bright probe flash (~16 or 33 ms) was presented, followed by 2 s post-flash time. A saccade-flash pair, along with pre- and post-saccade durations, formed one sequence (illustrated in detail below Sequence 2). The post-flash time of one sequence was the pre-flash time of the next sequence. A single trial consisted of 39 such sequences. In some sequences, no flash was presented after the saccade (saccade-only sequence). Here, the next saccade occurred 4 s after the previous saccade. The delay and the flash polarity were both randomized within a trial. In different trials, the saccades always remained the same but the order of delay and flash polarity changed. Each trial lasted for 2-4 minutes, and the number of trials needed depended on the number of conditions that were tested in an experiment. For example, to test 7 different probe-flash delays, 15 trials would be required: 7 trials for bright flashes, 7 trials for dark flashes, and 1 trial for no-flash condition. The pseudo-randomization within and across trials was designed in a way that a single condition (for example bright flash 150 ms after saccade onset), happened once after each of the 39 saccades.

**b.** Example trial of the luminance step paradigm. Here, the texture was replaced by a uniform gray background and saccades were represented by sudden increases or decreases in the background's luminance. In a single trial, 56 or 156 sequences occurred. In macaque RGC experiments, only 20 sequences occurred in a single trial. The luminance steps in a trial had different contrasts, spanning a range of -0.5 to +0.5 on the Michelson scale. Randomization was done similar to the saccade paradigm described in **a**. Each trial lasted for 3-7 minutes, and the number of trials needed depended on the number of conditions that were tested in an experiment.

**c.** Shortened luminance step paradigm used in cone imaging experiments. The luminance of a uniform 700 μm disc centered over the imaging field alternated between a bright and a dark intensity. The transitions caused positive- and negative-contrast luminance steps over the imaging field (+0.4 and -0.4 Michelson contrast). Dark (-0.33 Michelson contrast) or bright (+0.33) probe flashes (100 ms) occurred 50, 250 and 2000 ms after the step. A single trial consisted of 14 sequences: 2 contrast polarity x 2 flash polarity x 3 delays + 2 no flash conditions, and lasted ~45 s. Conditions were randomized within this trial.

**Figure S2**

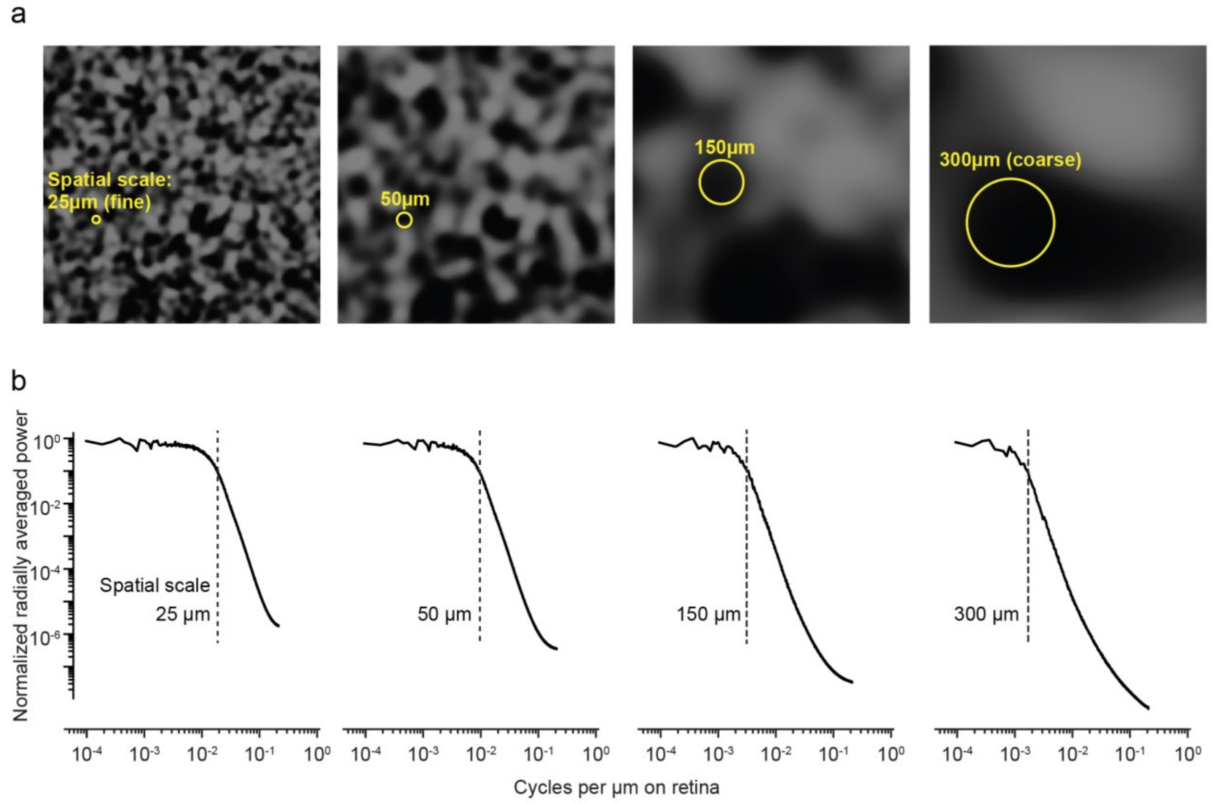


**Figure S2 Textured backgrounds.**

**a.** Textures were created by convolving random binary pixel images with a Gaussian blurring filter. We varied the $\sigma$ parameter of the Gaussian blurring filter to define a so-called spatial scale for the resulting texture (indicated as yellow circles). $\sigma$ was set to half the spatial scale value. The fine (25 μm) and coarse (150 - 300 μm) spatial scales were picked to result in dark or bright image blobs that approximated the sizes of bipolar cell and ganglion cell receptive fields respectively.

**b.** Radially-averaged power spectra for textures like in **a**, normalized to the maximum average power. Low-pass characteristics in all spatial scales were clear: less than 5% of the total average power was above the spatial frequency corresponding to the specific spatial scale of a given texture (vertical dashed lines).

**Figure S3**

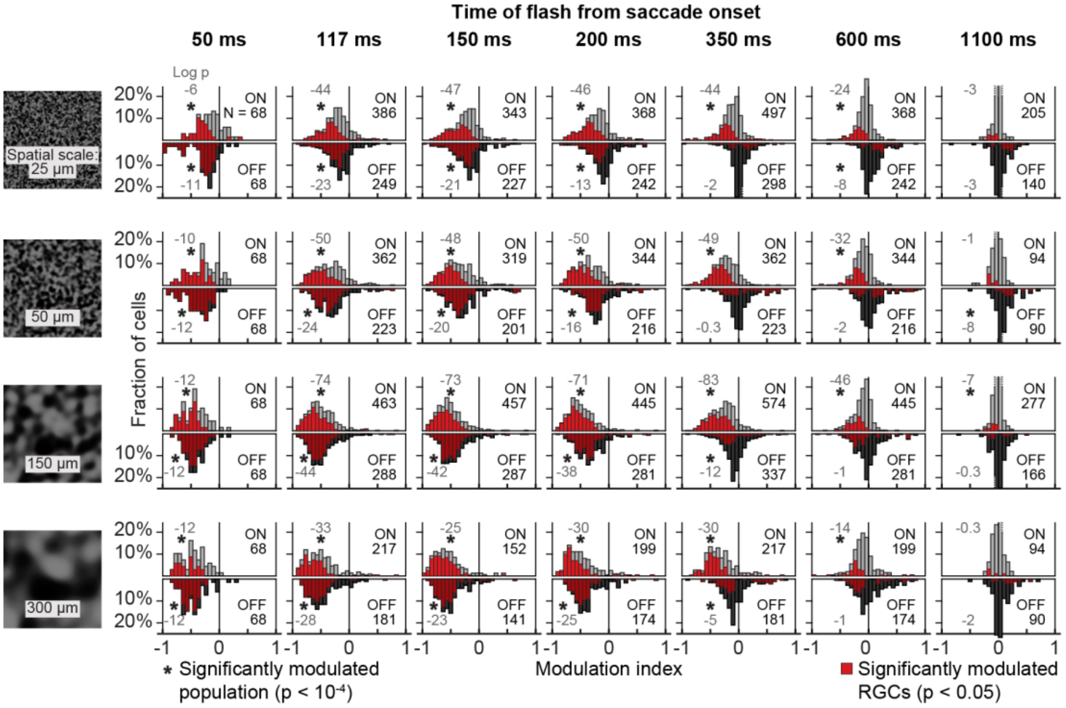


**Figure S3 Histograms of modulation indices of ON and OFF RGCs.** Histograms of modulation indices for all analyzed RGCs at different flash times relative to saccade onset (columns). Rows correspond to background textures of different spatial scales. Upright histograms (light bars) are for ON RGCs and inverted histograms (dark bars) are for OFF RGCs. These histograms show the underlying distribution for the mean modulation indices shown in Fig. 1e. Red bars indicate RGCs that are statistically significantly modulated (modulation index > 0 or < 0, $p < 0.05$, one-tailed sign test). Black numbers on the right in each panel indicate the number N of ON and OFF RGCs analyzed for that condition; gray numbers on the left in each panel are the logarithm (base 10) of the exact p-value (two-tailed Wilcoxon signed-rank test) to determine if the population median was shifted away from 0. Additionally, the asterisks show the significance level at $p < 0.001$.

**Figure S4**

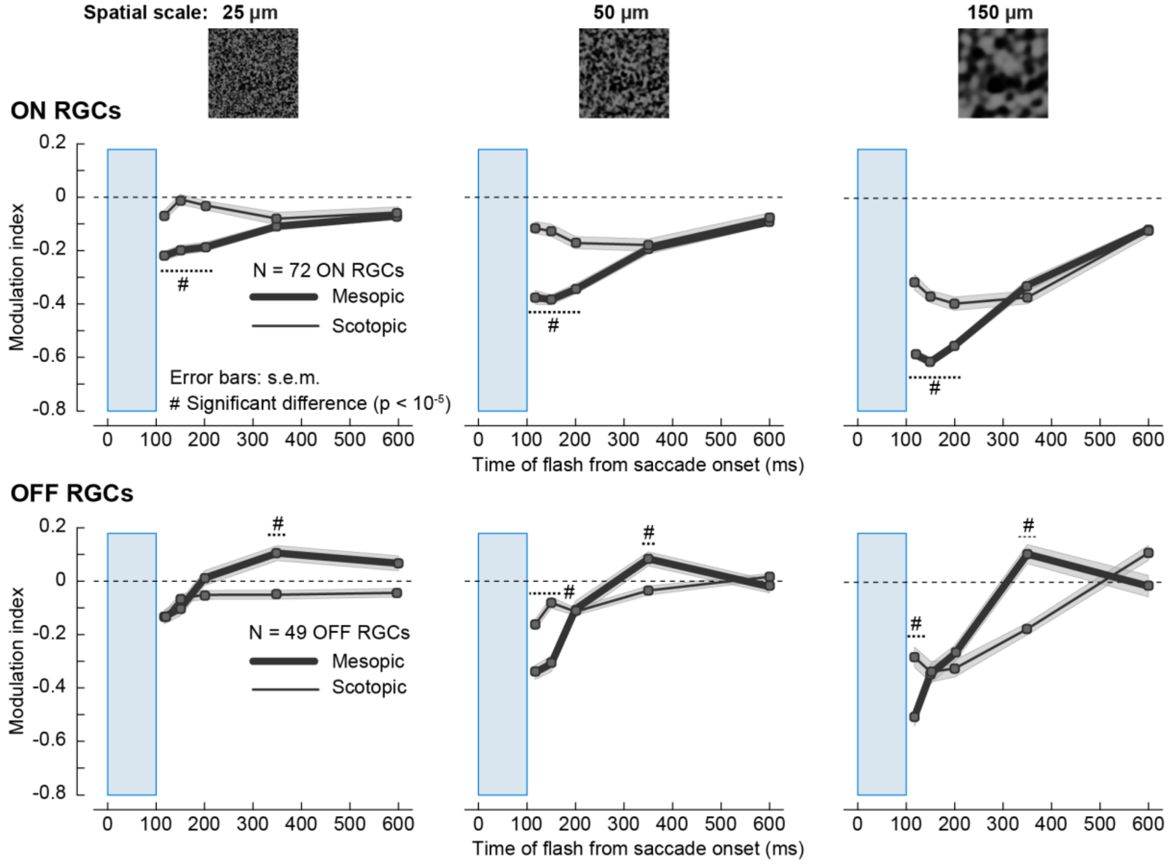


**Figure S4 Retinal saccadic suppression at scotopic light levels.** Population modulation index (mean ± s.e.m.) across ON (top; N = 72) and OFF (bottom; N = 49) RGCs at mesopic light levels (thick lines; mean absolute light intensity: 225 $R^{*}rod^{-1}s^{-1}$) and scotopic light levels (thin lines; mean absolute light intensity: 23 $R^{*}rod^{-1}s^{-1}$). Columns are for different spatial scales of the background texture. Mouse and pig RGC data shown in all other figures is from mesopic light levels. Suppression in both ON and OFF RGCs was weaker at scotopic light level. Blue window: timing of the saccade. Probe flashes were presented after saccade onset at 117, 150, 200, 350, 600 and 2100 ms. Hash symbols: statistically significant difference in modulation across mesopic and scotopic light levels ($p < 10^{-5}$, two-tailed Wilcoxon rank-sum test). Suppression at later time points (> 300 ms, presumably originating from the surround component, see Figs. 2, 3) was not different between mesopic and scotopic conditions in ON RGCs, and tended to be even slightly stronger in OFF RGCs. Suppression at early time points (< 250 ms, likely dominated by the central component, see Figs. 2, 3) was much less pronounced at scotopic conditions.

87 **Figure S5**

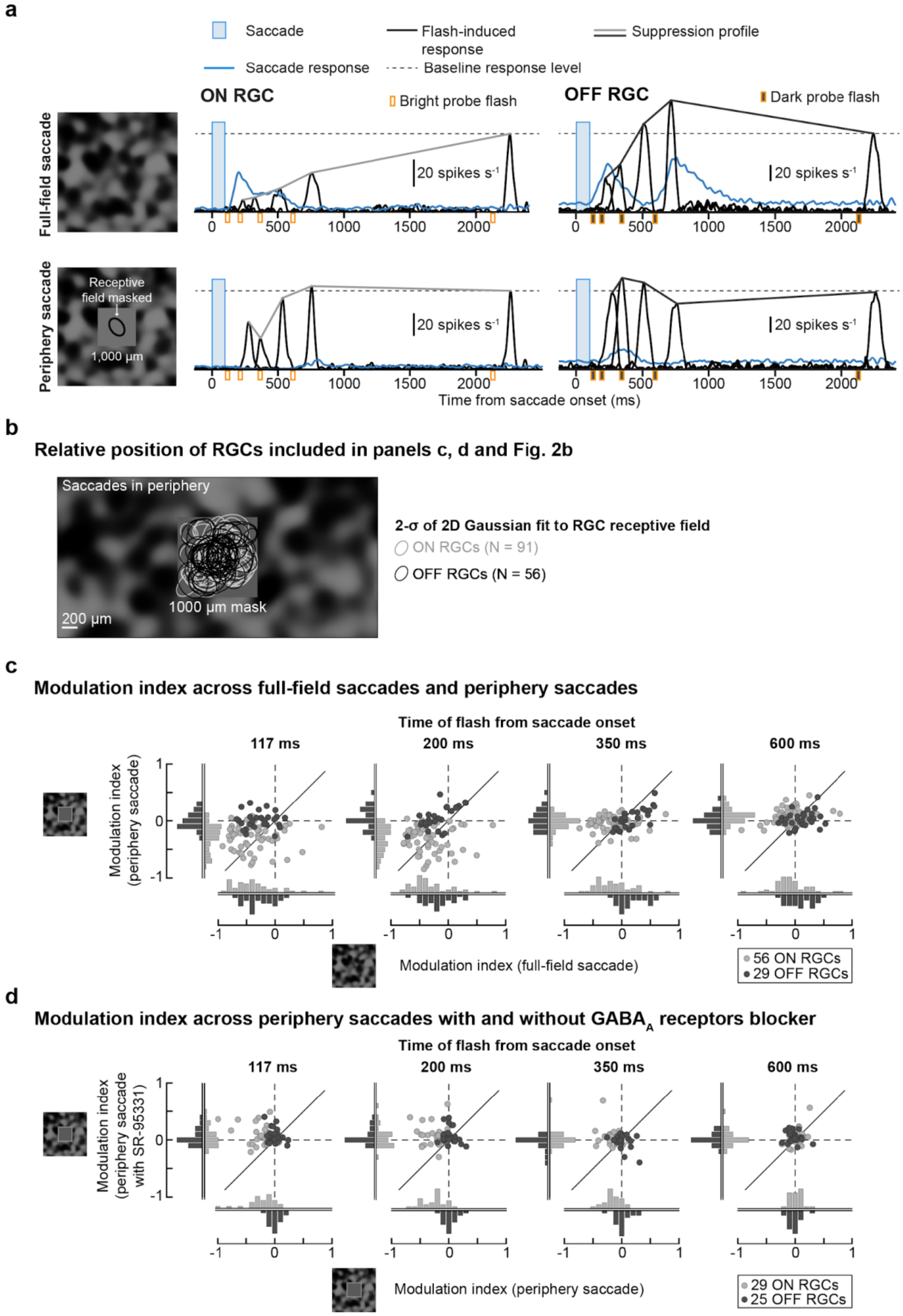


88
89
90

**Figure S5 Extended data figure for global component of suppression (Fig. 2a-b).**

**a**. Activity (firing rate) of example ON and OFF RGCs to full-field saccades (top; same paradigm as in Fig. 1; background texture here had a spatial scale of 300 μm) and during periphery saccades condition (bottom), where a 1000 x 1000 $\mu m^2$ gray square mask (intensity: mean luminance of texture) restricted saccades to RGC receptive field periphery (>2-$\sigma$ of the 2D Gaussian fit to the receptive field). Blue lines: average saccade-alone responses (N = 39 sequences). Black lines: average flash-induced responses (composite saccade and flash responses minus saccade-alone response). Blue windows: timing of saccades; orange markers: timing of the probe flashes; dashed lines: baseline response level (response to flash at 2100 ms). Periphery saccades revealed a global component of suppression in the ON RGC, which was weaker and short-lived (recovered by 350 ms) than full-field saccades. The OFF RGC was no longer suppressed with periphery saccades. Lines connecting the response peaks highlight the time courses of retinal saccadic suppression relative to baseline flash-induced responses.

**b.** Schematic showing position of the RGCs relative to the 1000 μm mask that were included in the analysis of **c**, **d**, Fig. 2b. These RGCs had at least 2-$\sigma$ of the 2D Gaussian fit to their receptive fields covered by the mask, shown here by bright and dark ellipses for ON (N = 91) and OFF (N = 56) RGCs, respectively. Saccades were restricted to outside of the mask, i.e. periphery of RGCs receptive fields.

**c.** Modulation indices of ON RGCs (light gray circles; N = 56) and OFF RGCs (dark gray circles; N = 29) across full-field saccades (x-axis) and periphery saccades (y-axis), at different flash times relative to saccade onset (columns). Oblique lines are the unity lines between the two conditions and the dashed lines correspond to zero modulation. Distribution of modulation indices for the two conditions are projected onto their respective axes (light gray bars for ON RGCs and dark gray bars for OFF RGCs). The modulation index of most OFF RGCs was close to 0 in the periphery saccades condition, for all flash times, suggesting they were not suppressed by the global component of suppression, originating from the receptive field periphery. ON RGCs were still suppressed, although they recovered by 350 ms, which was quicker than after full-field saccades.

**d.** Modulation indices of ON (N = 29) and OFF (N = 25) RGCs across periphery saccades without (x-axis) and with GABAA receptor blocker (y-axis; 5 μM SR-95531). With GABAA receptors blocked, modulation indices of ON RGCs were around 0, suggesting that the global component of suppression was mediated through GABAergic pathways. OFF RGCs remain unaffected.

#### Figure S6

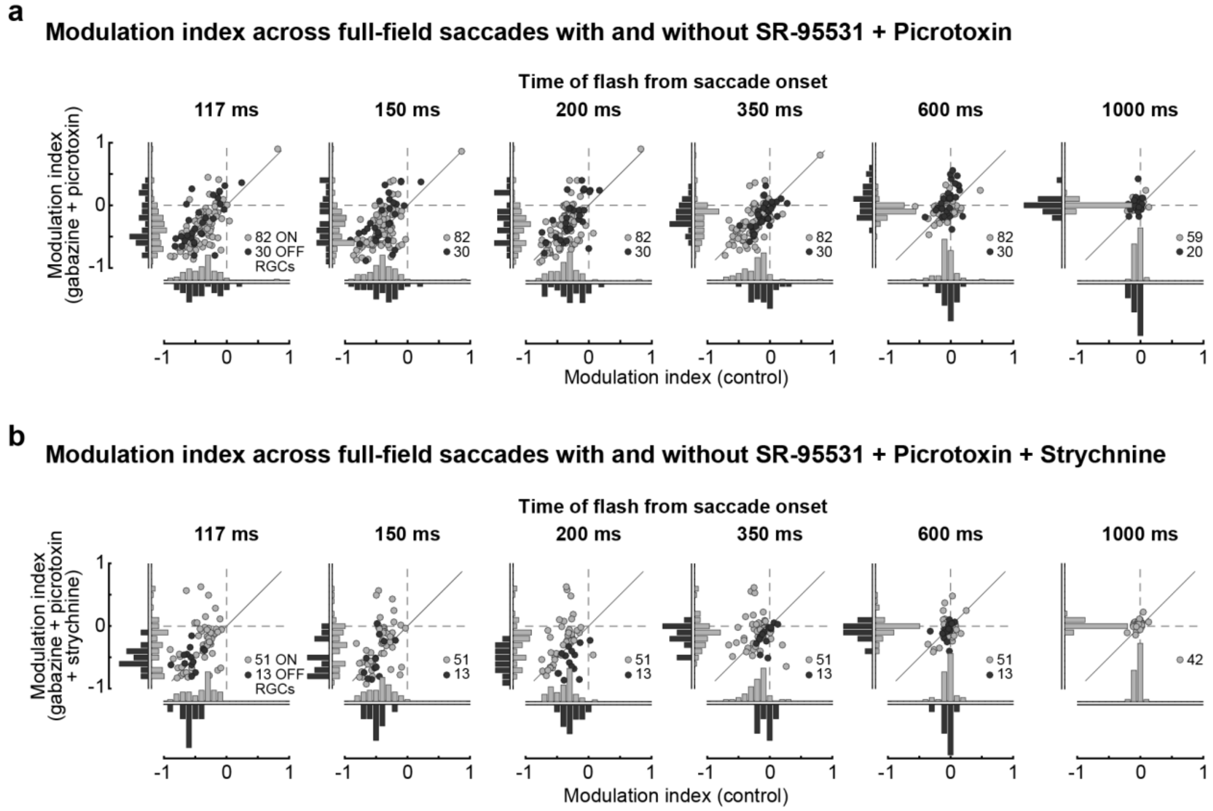


**Figure S6 Population data underlying pharmacology experiments of Fig. 2c, d.**
**a.** Modulation indices of ON RGCs (light gray circles; N = 59) and OFF RGCs (dark gray circles; N = 20) across full-field saccades in control conditions (x-axis) and with $GABA_{A,C}$ receptors blockers (y-axis; 5 μM SR-95531 + 100 μM Picrotoxin), at different flash times relative to saccade onset (columns). Oblique lines are the unity lines between the two conditions and the dashed lines correspond to zero modulation. Distribution of modulation indices for the two conditions are projected onto their respective axes (light gray bars for ON RGCs and dark gray bars for OFF RGCs). While the modulation index of individual cells showed small changes across the conditions, in general, the population was mostly concentrated around the unity line.
**b.** Same as in **a**, except that glycine receptors (1 μM Strychnine) were also blocked in addition to $GABA_{A,C}$ receptors. No OFF RGCs were recorded in experiments which included flashes at 1000 ms.

143 **Figure S7**

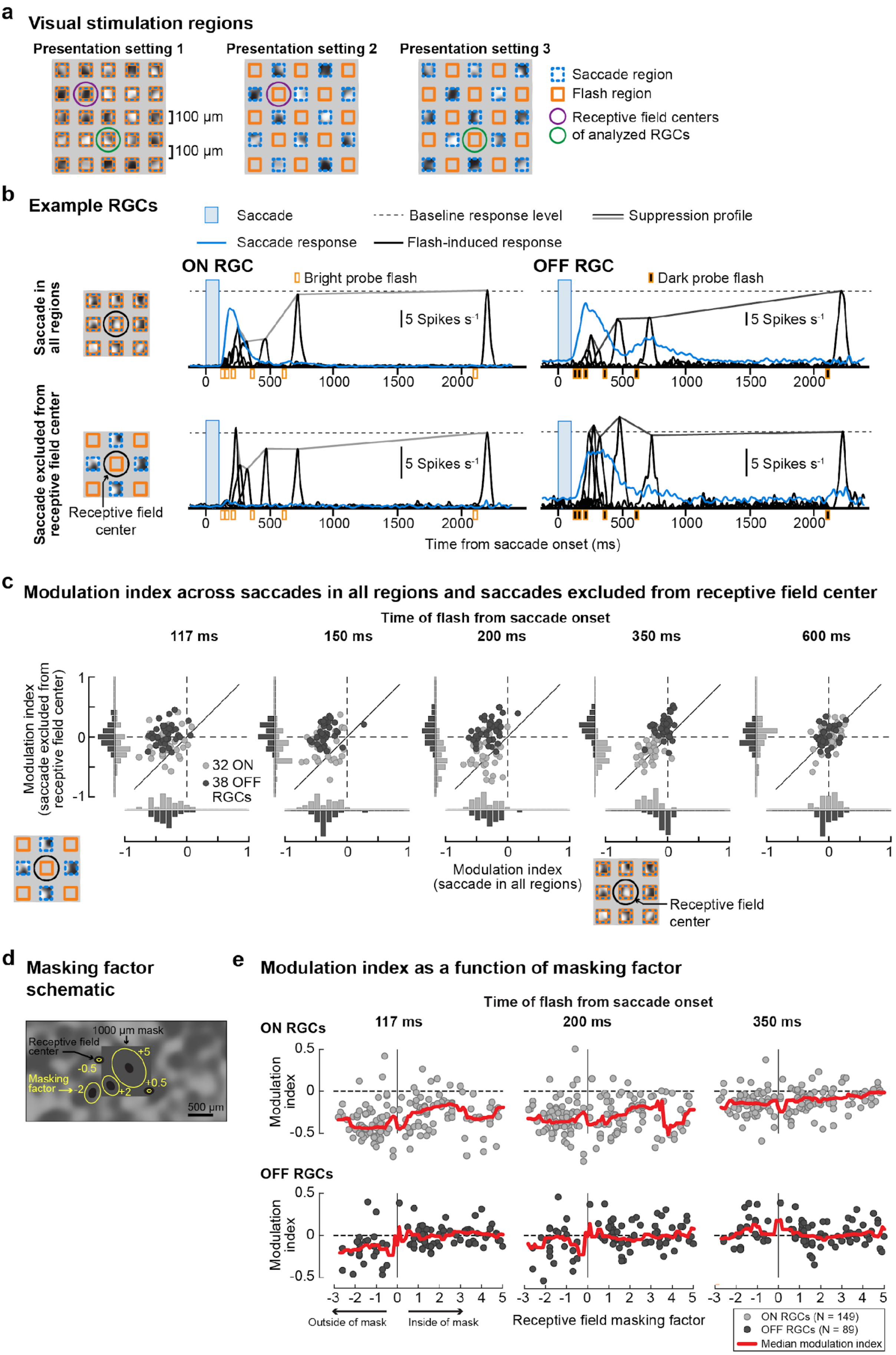


144

**Figure S7 Extended data figure for the local component of suppression (Fig. 2e)**

**a.** Schematic of the visual stimulation settings used in the checkerboard-mask paradigm. In setting 1, saccades and flashes were presented in all square regions of size 100 x 100 $\mu m^2$ separated by a gap of 100 $\mu m$. Throughout an experiment, the gap's intensity remained at mean luminance of the saccade background texture. In setting 2 and 3, saccades and flashes were presented in alternate sets of square regions. For quantifying modulation when saccades were excluded from the receptive field center, setting 2 was used for an RGC with position indicated by the purple circle and setting 3 was used for an RGC with position indicated by the blue circle.

**b.** Activity (firing rate) of example ON and OFF RGCs to the condition where saccades and flashes were presented in all regions (top), and where saccades and flashes were presented in alternate regions (bottom). In these experiments, we used a coarse background texture (150 $\mu m$ spatial scale). Blue lines: average saccade-alone responses (N = 39 sequences). Black lines: average flash-induced responses (composite saccade and flash responses minus saccade-alone response). Blue windows: timing of saccades; orange markers: timing of the probe flashes; dashed lines: baseline response level (response to flash at 2100 ms). Lines connecting the response peaks highlight the time courses of retinal saccadic suppression relative to baseline flash-induced responses.

**c.** Modulation indices of ON RGCs (light gray circles; N = 32) and OFF RGCs (dark gray circles; N = 38) across saccades and flashes presented in all square regions of a checkerboard mask (x-axis) and saccades and flashes presented in alternate regions of the checkerboard mask where saccades were excluded from the receptive field center of the analyzed RGC (y-axis), at different flash times relative to saccade onset (columns). Oblique lines are the unity lines between the two conditions and the dashed lines correspond to zero modulation. Distribution of modulation indices for the two conditions are projected onto their respective axes (light gray bars for ON RGCs and dark gray bars for OFF RGCs).

**d**. A masking factor was computed for each RGC recorded under the periphery saccades protocol (Fig. 2a). This factor was defined as the multiple of $\sigma$ of the 2D Gaussian fit of a cell's receptive field center (black filled ellipses) for which the ellipse just touched the mask boundary (yellow ellipses). Positive masking factor: cells with receptive field centers within the mask; negative factor: cells with receptive field centers outside the mask. The magnitude of the factor increased with distance of the receptive field from the edge of the mask. Only RGCs with masking factors > 2 were included in the analysis of global component of suppression (Fig. 2b, S5).

**e.** Modulation indices of ON RGCs (top; light circles; N = 149) and OFF RGCs (bottom; dark circles; N = 89) plotted as a function of receptive field masking factor at different flash times relative to saccade onset (columns). Red lines indicate the running median modulation index (Methods). In OFF RGCs with more than half of their receptive fields covered by the mask (masking factor > 0), only weak or no suppression of flash responses was observed. However, when more than half of the receptive field of OFF RGCs was exposed to the saccade, even marginally (masking factors between 0 and -1), flash responses were suppressed. This confirms the findings of Fig. 2f and S7c, that there is a narrow spatial window centered on the receptive field center of OFF

RGCs, which, when stimulated with a saccade and a subsequent flash, suppresses the response to the flash.

**Figure S8**

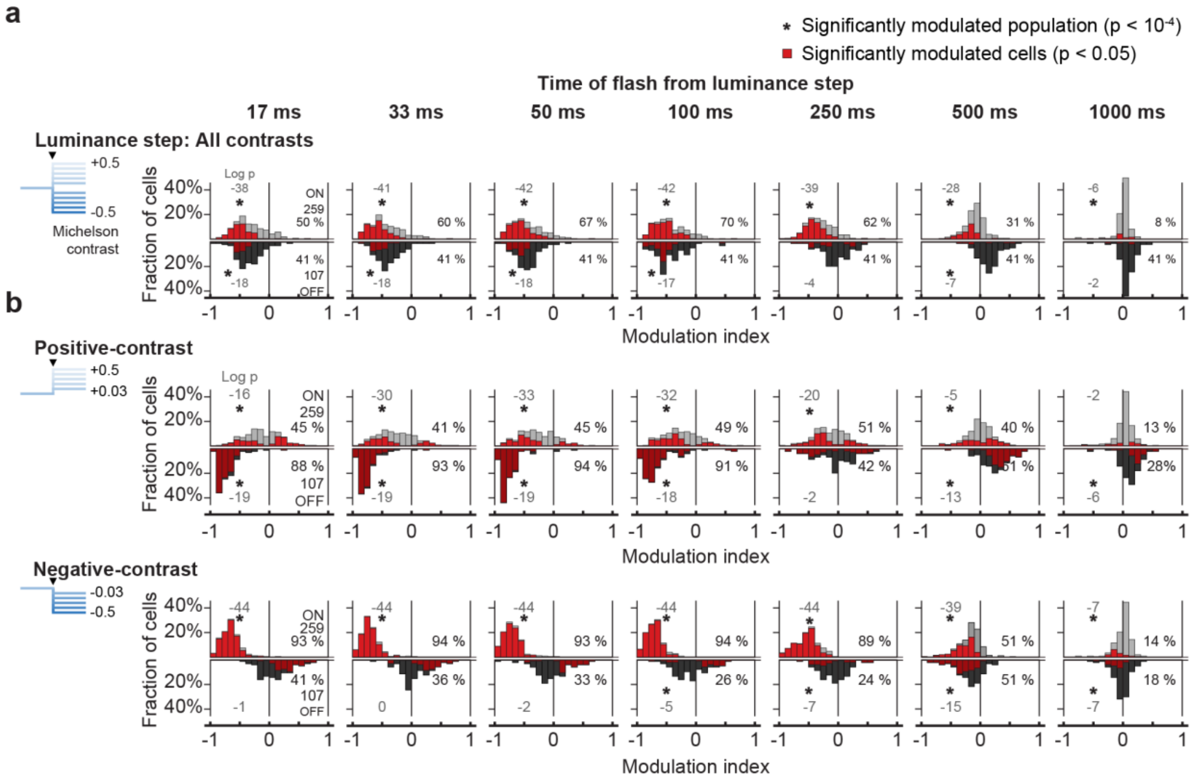


**Figure S8 Population data underlying luminance step experiments of Fig. 4.** Histograms of modulation indices for all analyzed RGCs. Modulation index for each RGC was based on responses averaged across all luminance steps collectively (**a**; -0.5 to +0.5 Michelson contrast; N = 56 or 156 sequences), or across all positive-contrast (N = 28 or 78 sequences) and all negative-contrast (N = 28 or 78 sequences) luminance steps separately (**b**). Upright histograms (light gray bars) are for ON RGCs (N = 259) and inverted histograms (dark gray) are for OFF RGCs (N = 107). Red bars indicate the RGCs with statistically significant modulation (modulation index < 0 or > 0, p < 0.05, one-tailed sign test). Columns show modulation at different flash times relative to luminance steps. These histograms show the underlying distribution for the mean population modulation indices shown in Fig. 4. The percentage values on the right in each panel indicate the percentage of RGCs significantly modulated in that condition. Gray value on the left in each panel is the logarithm (base 10) of the exact p-value (two-tailed Wilcoxon signed-rank test) to determine if the population median was shifted away from 0. Additionally, the asterisks indicate significance level p < 0.0001.

212

### Figure S9

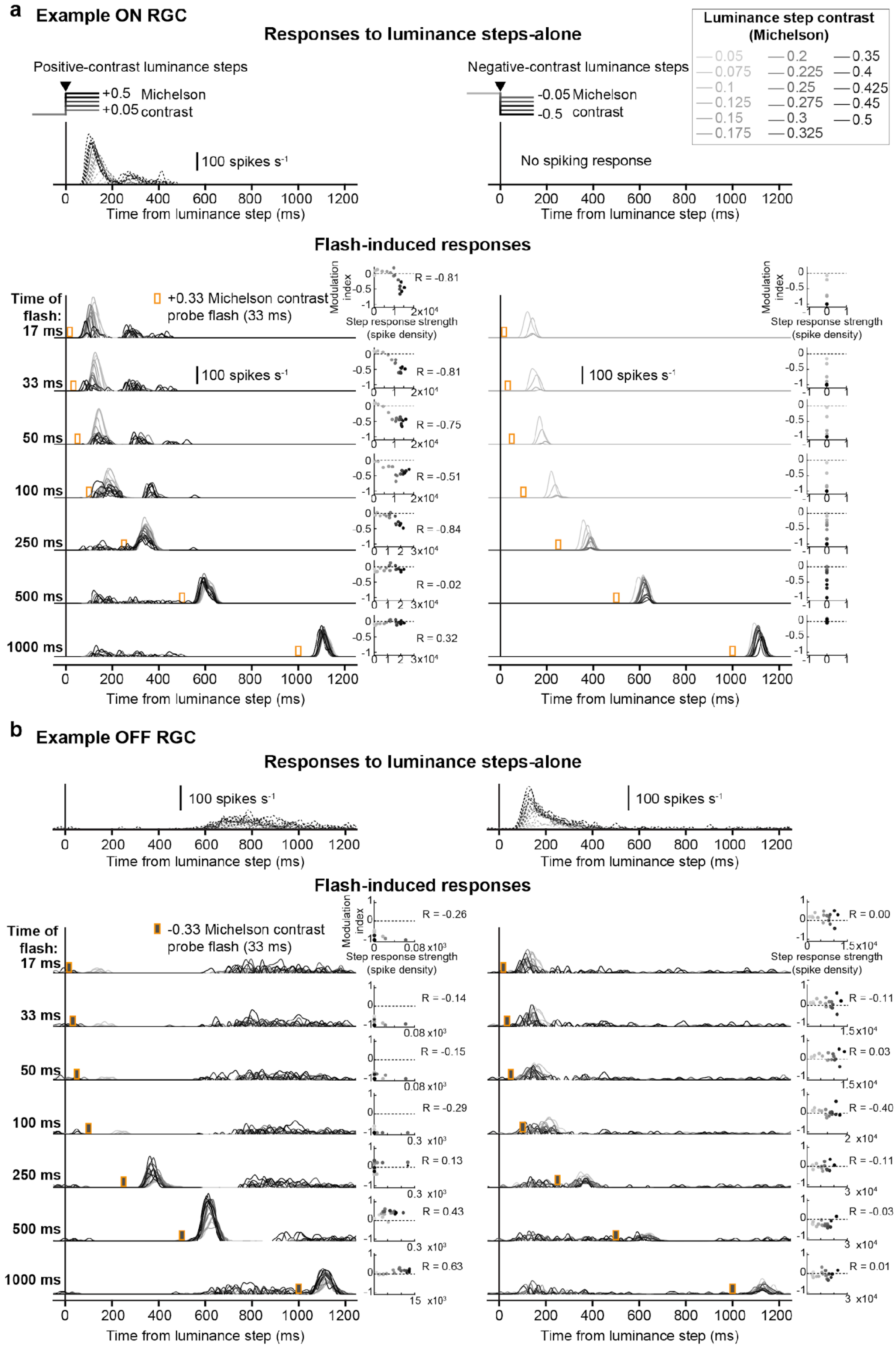


213

**Figure S9 Example cells showing suppression of flash responses following luminance steps.**

**a.** Average activity of an example ON RGC to positive-contrast (left column) and negative contrast (right column) luminance steps. Top row: luminance steps alone (dotted lines). Other rows: flash-induced responses (solid lines; different rows represent flashes presented with different delays), obtained by subtracting responses to the isolated luminance step from responses to the composite luminance step and bright flash stimulus. Responses to the different luminance step → flash sequences (N = 56 or 156; Fig. S1b) were first binned according to the contrast induced by the luminance step (17 bins; 0.05 to 0.5 absolute Michelson contrast range). Responses were then averaged within each bin, as shown here. Darker lines represent responses after stronger-contrast luminance steps. Orange bars indicate time of probe flash after the step (+0.33 Michelson contrast; 33 ms long). After positive-contrast luminance steps (left column), the RGC responded relatively strongly to flashes presented after weak luminance steps (lighter lines). However, responses to flashes presented 17-100 ms after strong luminance steps were suppressed, as indicated by the reduced amplitude or even absence of dark traces in these panels. The scatter plots towards the right show the dependency of flash-induced response modulation (y-axis; modulation index, Methods) on the luminance-step response strength (x-axis; Methods). In general, stronger positive-contrast luminance steps (darker circles) induced stronger responses to luminance-steps themselves (darker dots are further right), and also caused stronger suppression of flash-induced responses for flashes presented 17-250 ms after the step (darker dots are further down). This monotonic relation between flash response modulation index and luminance step response strength was quantified by calculating the Spearman correlation, R, which we refer to as the association index (AI). The larger its magnitude, the stronger is the monotonic relation between the two quantities. A negative AI value indicates a negative monotonic relation such that stronger luminance step responses are associated with decreasing (more negative) modulation indices (i.e. weaker flash responses → stronger suppression); a positive AI value indicates that stronger luminance step responses are associated with increasing modulation indices (i.e. stronger flash responses → less suppression or even enhancement). The example ON RGC showed no spiking responses to negative-contrast luminance steps (right column), and therefore the association index was undefined. Nonetheless, stronger-contrast luminance steps caused stronger suppression, noted by the absence of dark response lines for flashes presented up to 100 ms after the step.

**b.** Same as in **a**, but for an example OFF RGC where probe flashes were dark (-0.33 Michelson contrast). Dark flash responses were strongly suppressed after a positive-contrast luminance step (17-100 ms; left column), irrespective of the step contrast. Responses were enhanced around 500 ms with stronger contrasts inducing stronger enhancement (note the positive AI). Responses to flashes presented after a negative-contrast luminance step (right column) were not suppressed.

**Figure S10**

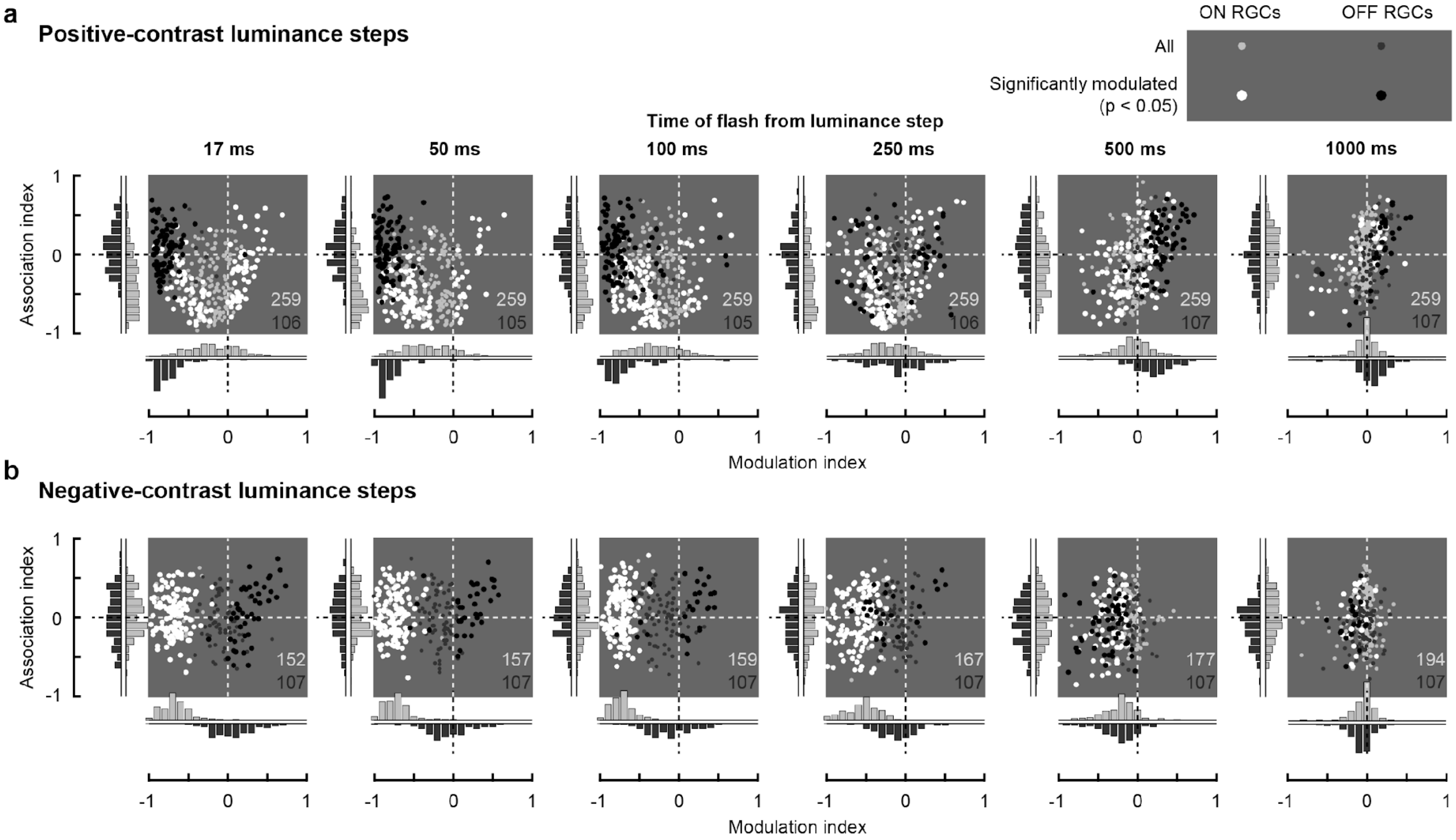


**Figure S10 Indication of suppression through a saturation-like mechanism in ON RGCs.** Scatter plots showing association index (AI, y-axis) of ON and OFF RGCs as a function of their modulation index (x-axis), at different flash times after positive-contrast **(a)** and negative-contrast luminance steps **(b)**. See Legend of Fig. S9 and Methods for an explanation of the association index. These RGCs are a subset of cells in Fig. 4, S8. Light gray and dark gray circles are all ON and OFF RGCs respectively; white and black circles are the significantly modulated ON and OFF RGCs ($p < 0.05$, one-tailed sign test) respectively. Dashed lines represent no modulation (vertical) and no association (horizontal). Distribution of the indices for all RGCs are projected onto the relevant x- and y-axes (light gray bars for ON RGCs and Dark gray bars for OFF RGCs). After a positive-contrast luminance step (**a**), many ON RGCs are suppressed (negative modulation index, although not as strongly as after a negative-contrast step) and have a negative association index (see example ON RGC in Fig. S9a). Such ON RGCs are located in the lower-left quadrants of the scatter plots in **a**. In these RGCs, suppression of flash responses increased monotonically (i.e. the modulation index decreased), with increasing response to the preceding luminance steps. This is consistent with an adaptation/saturation mechanism, where the response to a positive-contrast luminance step might strongly activate ON RGCs, so that the response to a subsequent probe flash would drive the cells into adaptation or saturation, effectively resulting in suppressed flash responses. Note that there are no OFF RGCs with similar properties, as, in general, they were not suppressed by negative-contrast luminance steps; they would occupy the lower left quadrant in the scatter plots in **b**.

#### Figure S11

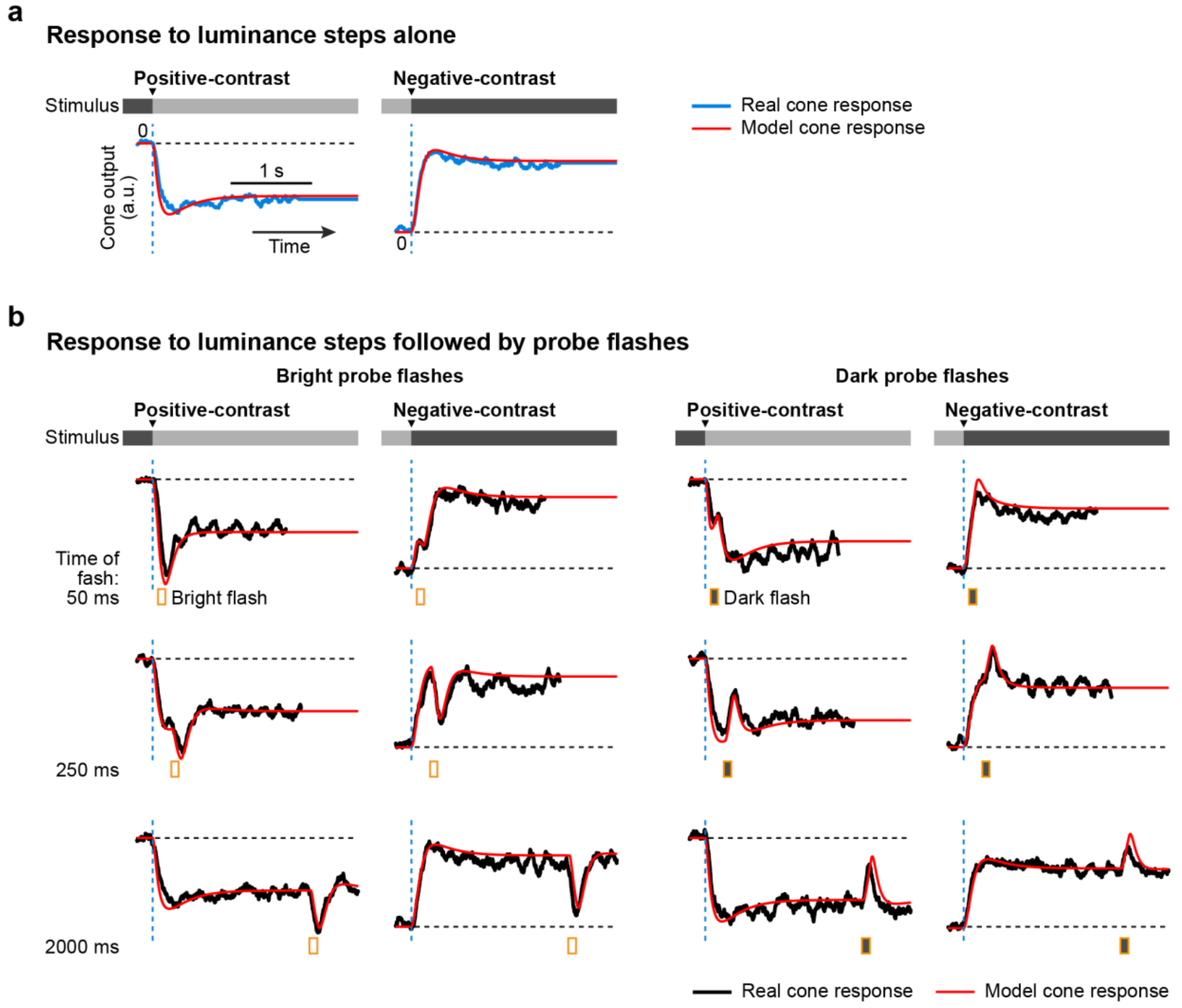


**Figure S11 Model fit to cone responses of Fig. 5.**

**a,b.** Cone responses to positive-contrast and negative-contrast (+0.4 and -0.4 on Michelson scale) luminance steps alone **(a)** and to luminance steps followed by probe flashes at 17, 250 and 2000 ms **(b)**. Probe flashes were either bright or dark (+0.33 or -0.33 Michelson contrast respectively; 100 ms long). Orange markers show the timing of probe flashes. Real cone responses (blue/black; normalized ΔF/F of the iGluSnFR indicator signal; same as Fig. 5a-b) and model cone responses (red) are overlaid. Dashed blue lines: timing of luminance step; horizontal dashed line: cone level prior to the luminance step.

#### Figure S12

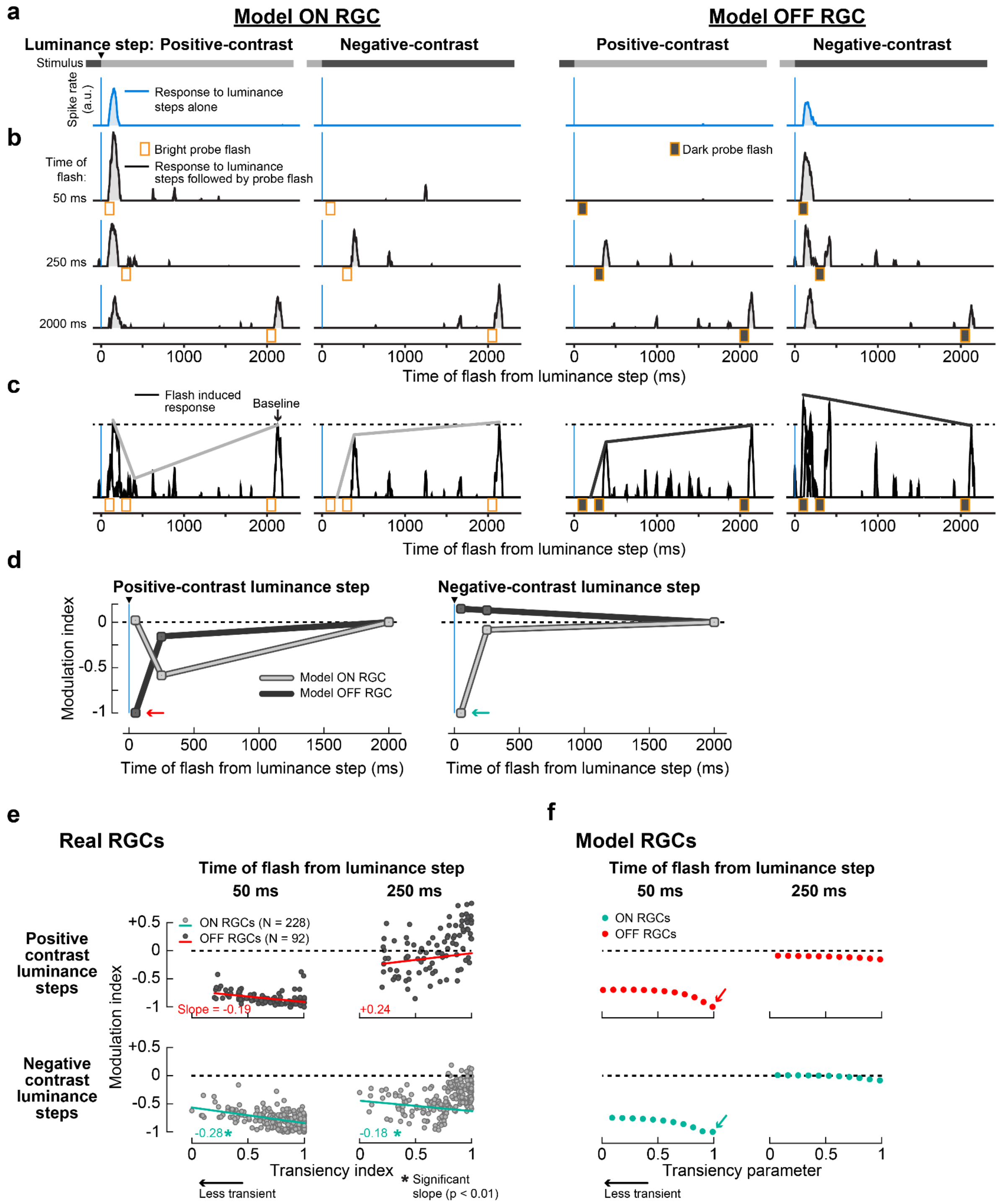


**Figure S12 Model RGC responses based on real cone data of Fig. 5.**
**a,b.** Spiking response of model ON (columns 1-2) and OFF (columns 3-4) RGCs to luminance steps alone (**a**, blue) and to luminance steps followed by probe flashes (**b**, black). First column in each cell type: responses following a positive-contrast luminance step; second column: responses following a negative-contrast luminance step. Instead of model cone output like in Fig. 6, we used acquired cone responses (Fig. 5) to RGC responses. Horizontal intensity bar below each trace illustrates the underlying visual stimuli. Vertical blue lines: timing of luminance step; orange bars: timing of probe flashes.

**c.** Flash-induced responses, after subtracting **a** from **b**, overlaid to show the modulation of probe flash responses at different times (analogous to real RGCs in Fig. 1d). Response to flash presented 50 ms after the step was strongly suppressed in ON RGC when presented after the negative-contrast step and strongly suppressed in OFF RGC when presented after the positive-contrast step, consistent with the cross-over style of suppression observed in real RGC data of Fig. 4, or in RGC models of Fig. 6.
**d.** Modulation indices for probe flashes in ON (light gray) and OFF model RGCs (dark gray), following negative-contrast (left panel) and positive-contrast (right panel) luminance steps. Cyan and red arrows highlight the suppression of opposite-contrast flashes at 50 ms in ON and OFF RGCs, respectively.
**e.** Replica of Fig. 6f but with relevant time points only. Modulation indices of real ON RGCs (light gray circles; N = 228; cyan line: linear regression fit) and OFF RGCs (dark gray circles; N = 92; red line: linear regression fit) plotted as a function of RGC transiency index (Methods). These RGCs are a subset of the population data shown in Fig. 4b for which we could compute a transiency index. Columns correspond to flashes presented at different times after positive-contrast (top row) and negative-contrast (bottom row) luminance steps. Suppression was stronger in more transient ON and OFF RGCs after negative- and positive-contrast steps respectively, indicated by the negative slope of the linear fits. Numbers in each panel indicate the slope of the fits and asterisk symbol indicates statistically significant slope (slope $\neq 0$, $p < 0.01$, two-tailed t-test).
**f.** Modulation indices of model ON (cyan) and OFF (red) RGCs plotted as a function of the model's filter transiency parameter. Individual panels correspond to different flash times (chosen from real-RGC recordings, Fig. 4) after positive-contrast (top row) and negative-contrast (bottom row) luminance steps. In **a-d**, the filter transiency parameter was set to 1. Arrows highlight the same data as in **d**.

**Figure S13**

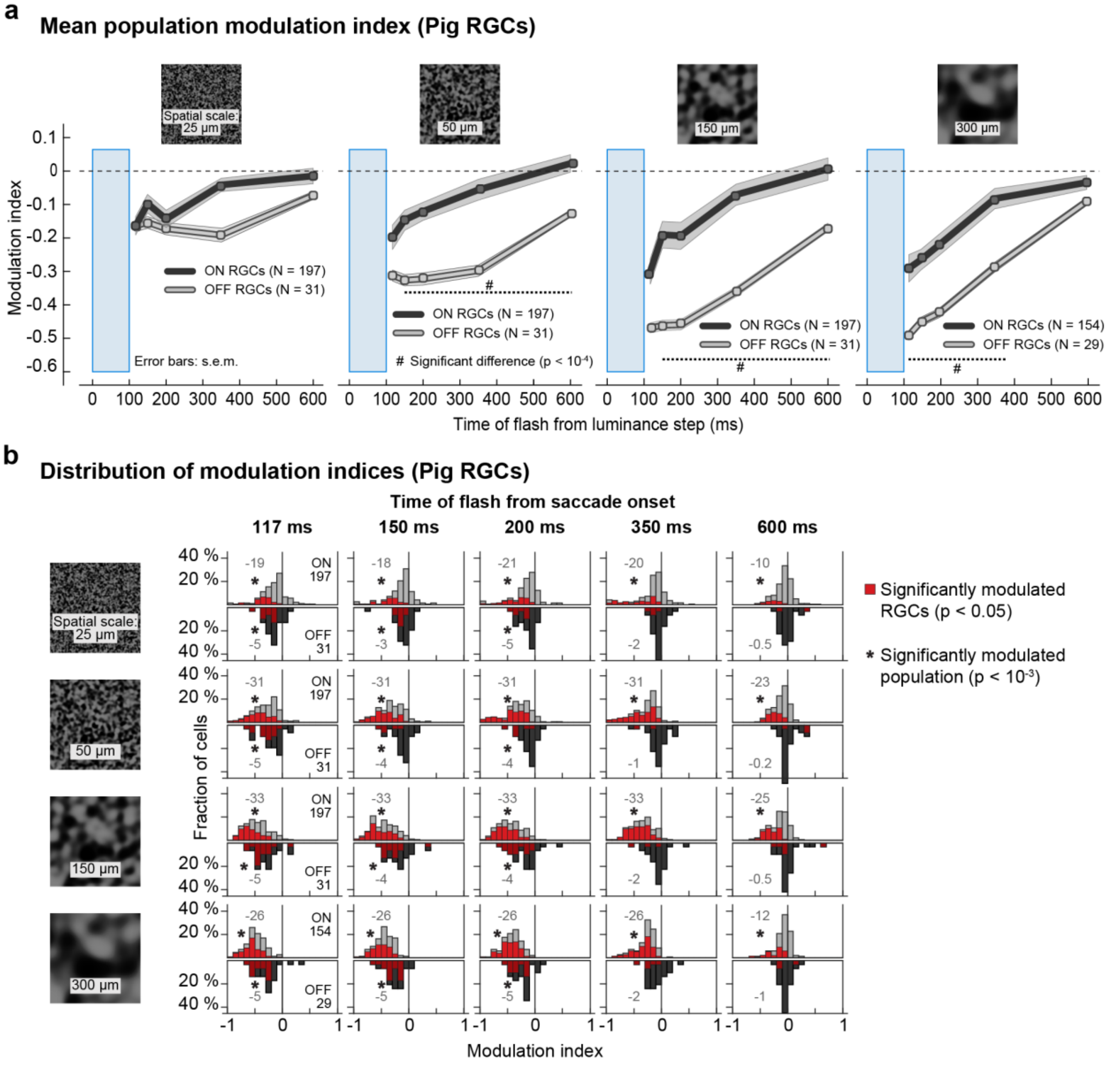


**Figure S13 Saccadic suppression in pig retina.**

**a.** Population modulation index (mean ± s.e.m.) across ON (light gray) and OFF (dark gray) RGCs, for different spatial scales of the background texture (columns). OFF RGCs recovered from suppression by 350 ms whereas ON RGCs recovered by 1 s. Suppression in both ON and OFF RGCs increased with the spatial scale of the background texture. These observations were consistent with observations from mouse RGCs (Fig. 1e, S3). Blue windows: timing of saccades. Probe flashes were presented at 117, 150, 200, 350, 600, and 2100 (baseline) ms after saccade onset. Population modulation index was based on average across 197 ON and 31 OFF RGCs for spatial scales 25, 50, 150 µm; and 154 ON and 29 OFF RGCs for 300 µm. Hash symbols: significant difference in modulation between ON and OFF RGCs ($p < 10^{-4}$, two-tailed Wilcoxon rank-sum test).

**b.** Histograms of modulation index for all analyzed pig RGCs at different flash times relative to saccade onset (columns). Rows are for background textures with different spatial scales. Upright histograms (light gray bars) are for ON RGCs and inverted histograms (dark gray) are for OFF RGCs. Red bars highlight the RGCs with modulation index significantly different from 0 ($p < 0.05$, one-tailed sign test). Black numbers in each panel of the first column indicate the number of RGCs analyzed for that background texture. Gray numbers on the left in each panel show the logarithm (base 10) of the exact p-value (two-tailed Wilcoxon signed-rank test) to determine if the population median was shifted away from 0. Additionally, the asterisks indicate the significance at $p < 0.001$.
